# Depletion of senescent cells improves functional recovery after spinal cord injury

**DOI:** 10.1101/2020.10.28.358762

**Authors:** Diogo Paramos-de-Carvalho, Isaura Martins, Ana Margarida Cristóvão, Ana Filipa Dias, Dalila Neves-Silva, Telmo Pereira, Diana Chapela, Ana Farinho, António Jacinto, Leonor Saúde

**Author notes:** Corresponding author: Leonor Saúde, Instituto de Medicina Molecular, Faculdade de Medicina da Universidade de Lisboa, Avenida Professor Egas Moniz, 1649-028 Lisboa, Portugal. These two authors ontributed equally to this work (Diogo Paramos-de-Carvalho, Isaura Martins).

## Abstract

Persistent senescent cells (SCs) are known to underlie ageing-related chronic disorders, but it is now recognized that SCs may be at the center of tissue remodeling events, namely during development or organ repair. Here we show that two distinct senescence profiles are induced in the context of a spinal cord injury between the regenerating zebrafish and the non-regenerating mouse. While induced-SCs in the zebrafish are progressively cleared out, they accumulate over time in mice. Depletion of SCs in spinal cord injured mice, with different senolytic drugs, improved locomotor, sensory and bladder functions. This functional recovery is associated with improved myelin sparing, reduced fibrotic scar, attenuated inflammation and increased axonal growth. Targeting SCs is a promising therapeutic strategy not only for spinal cord injuries but potentially for other organs that lack regenerative competence.

## INTRODUCTION

A spinal cord injury is a major cause of disability in humans and other mammals, often leading to permanent loss of locomotor and sensory functions. This type of traumatic lesion is defined by three biological features: a lesion core or fibrotic scar with no viable neural tissue; an astrocytic scar around the lesion core; and a surrounding area of spared neural tissue with limited function, which may exhibit some functional plasticity (O’Shea, Burda and Sofroniew, 2017). Although the lesion scar provides structural support, it also creates an inhibitory microenvironment for the regrowth of severed axons, thus preventing re-enervation of the original targets (Jared M. Cregg *et al.*, 2014). A spinal cord injury is further defined as an inflammatory condition mediated by activated astrocytes and infiltrating macrophages that remain in the spinal cord indefinitely (Donnelly and Popovich, 2008). Immediately after the injury, the blood-spinal cord barrier is disrupted and, although it gradually recovers, it remains compromised for a long period (Whetstone *et al.*, 2003). This facilitates the extravasation of immune cells contributing to the establishment of a chronic inflammatory state (Beck *et al.*, 2010).

In contrast to mammals, the zebrafish spinal cord has the remarkable capacity to recover motor and sensory functions after injury. This regenerative ability seems to stem from the supportive microenvironment where there is no formation of a glial or fibrotic scar and inflammation is dynamically controlled by macrophages (Tsarouchas *et al.*, 2018), allowing neurogenesis and regrowth of severed axons (Becker *et al.*, 1997; Vajn *et al.*, 2013).

While considerable knowledge was achieved on the biological processes that occur after a spinal cord injury in mammals and regenerative species, small progress was obtained on therapeutic options, suggesting that other cellular players might be relevant following an injury. Senescence is a cellular concept traditionally seen as an irreversible cell-cycle arrest response related to aging (van Deursen, 2014; Herranz and Gil, 2018; Gorgoulis *et al.*, 2019). Studies in recent years changed the way we perceive cellular senescence, placing it at the center of tissue remodeling in disease settings by limiting fibrosis, namely in wound healing (Jun and Lau, 2010; Demaria *et al.*, 2014), damaged livers (Krizhanovsky *et al.*, 2008; Kong *et al.*, 2012) and infarcted hearts (Meyer *et al.*, 2016). In regenerative models, such as salamander limbs, zebrafish hearts and fins and neonatal mouse hearts, a burst of transient senescent cells (SCs) was shown to be induced after an injury (Yun, Davaapil and Brockes, 2015; Da Silva-Álvarez *et al.*, 2019; Sarig *et al.*, 2019). These cells were shown to be efficiently cleared from the tissues as regeneration progressed possibly by macrophages (Yun, Davaapil and Brockes, 2015). Remarkably, if this initial senescence is eliminated, zebrafish fin regeneration is impaired (Da Silva-Álvarez *et al.*, 2019), suggesting that a transient accumulation of SCs appears to have beneficial functions. Persistent senescence, on the other hand, is detrimental for tissue and organ function in ageing and aged-related diseases, such as atherosclerosis, osteoporosis, diabetes and neurodegeneration (Calcinotto *et al.*, 2019). Key to their various roles is the fact that SCs secrete a plethora of factors known as senescence-associated secretory phenotype (SASP). It is through their SASP that SCs communicate with neighboring cells and modulate the tissue microenvironment, thus exerting most of their physiological effects (Acosta *et al.*, 2013; Calcinotto *et al.*, 2019). Importantly, the SASP mediates paracrine senescence, a process where senescent cells induce neighboring cells to undergo senescence (Herranz and Gil, 2018). It is becoming accepted that the beneficial versus detrimental effects of the SASP depend not only on its composition and stage of senescence progression, but also on the cell type affected and the stressor/injury type (Herranz and Gil, 2018).

Considering that persistent senescence was shown to be harmful, we hypothesized that accumulation of SCs contributes to the failure of spinal cord regeneration observed in mammals. In agreement with this hypothesis we have shown that SCs are induced after spinal cord injury in both zebrafish and mice. Whereas induced SCs in the zebrafish spinal cord progressively decrease, in mice these cells increase over time. Strikingly, the pharmacological depletion of SCs during the acute post-injury phase in mice seems to attenuate the secondary damage and maximize the extent of spared tissue, by decreasing inflammation burden, scar extension and demyelization, leading to a better functional outcome. Therefore, our data support the potential use of therapeutics targeting SCs to promote neuroprotection in the context of spinal cord injuries.

## RESULTS

### Zebrafish and mice exhibit distinct senescence profiles after spinal cord injury

A transient senescent profile was recently described in several regenerating organs after an injury (Yun, Davaapil and Brockes, 2015; Da Silva-Álvarez *et al.*, 2019; Sarig *et al.*, 2019) but the senescence profile in injured non-regenerating organs was never characterized.

To determine if senescence is induced following a spinal cord lesion in zebrafish and in mouse, we used the gold standard method to identify SCs – the senescence-associated β-galactosidase (SA-β-gal) assay (Itahana, Campisi and Dimri, 2007). In the regenerative zebrafish, SA-β-gal^+^ cells were found mainly in the spinal cord grey matter of Sham-injured animals (Fig. 1A). Strikingly, upon an injury, these cells were induced at the lesion periphery, reaching a peak at 15 days post-injury (dpi), when they double in number, and then returning to basal levels at 60 dpi (Fig. 1A). In scaring non-regenerative mice, SA-β-gal^+^ cells were also detected mainly in the spinal cord grey matter of Sham-injured animals and induced at the lesion periphery upon injury (Fig. 1B). However, in clear contrast to the zebrafish, these SA-β-gal^+^ cells did not return to basal levels and instead accumulated over time, reaching a four-fold increase at 60 dpi, when compared with Sham animals (compare Fig. 1A and Fig. 1B). Importantly, we confirmed that these profiles were injury-driven and not age-dependent by showing that the number of SA-β-gal^+^ cells remained unchanged in the spinal cord of Sham-injured animals of different ages, spanning all experimental time-points of this study (**Fig. S1**). We further confirmed by immunofluorescence the association of SA-β-gal^+^ cells with several senescence-associated biomarkers (Calcinotto *et al.*, 2019). In zebrafish, we were able to show that SA-β-gal^+^ cells co-localized with the cell cycle arrest marker p21^CIP1^ (Fig. 1C) and were devoid of the proliferation marker BrdU (Fig. 1D). In mice, SA-β-gal^+^ cells co-localized with the cell cycle arrest marker p16^INK4a^ (Fig. 1E) and the DNA damage marker γH2AX (Fig. 1F). In zebrafish and in mice, some SA-β-gal^+^ cells exhibited a clear enlarged morphology, another hallmark of SCs (Narita *et al.*, 2003). These results reveal two clearly distinct senescence profiles in an injured spinal cord, a transient profile observed in regenerating zebrafish and a persistent one observed in non-regenerating mice.

**Fig. 1.**
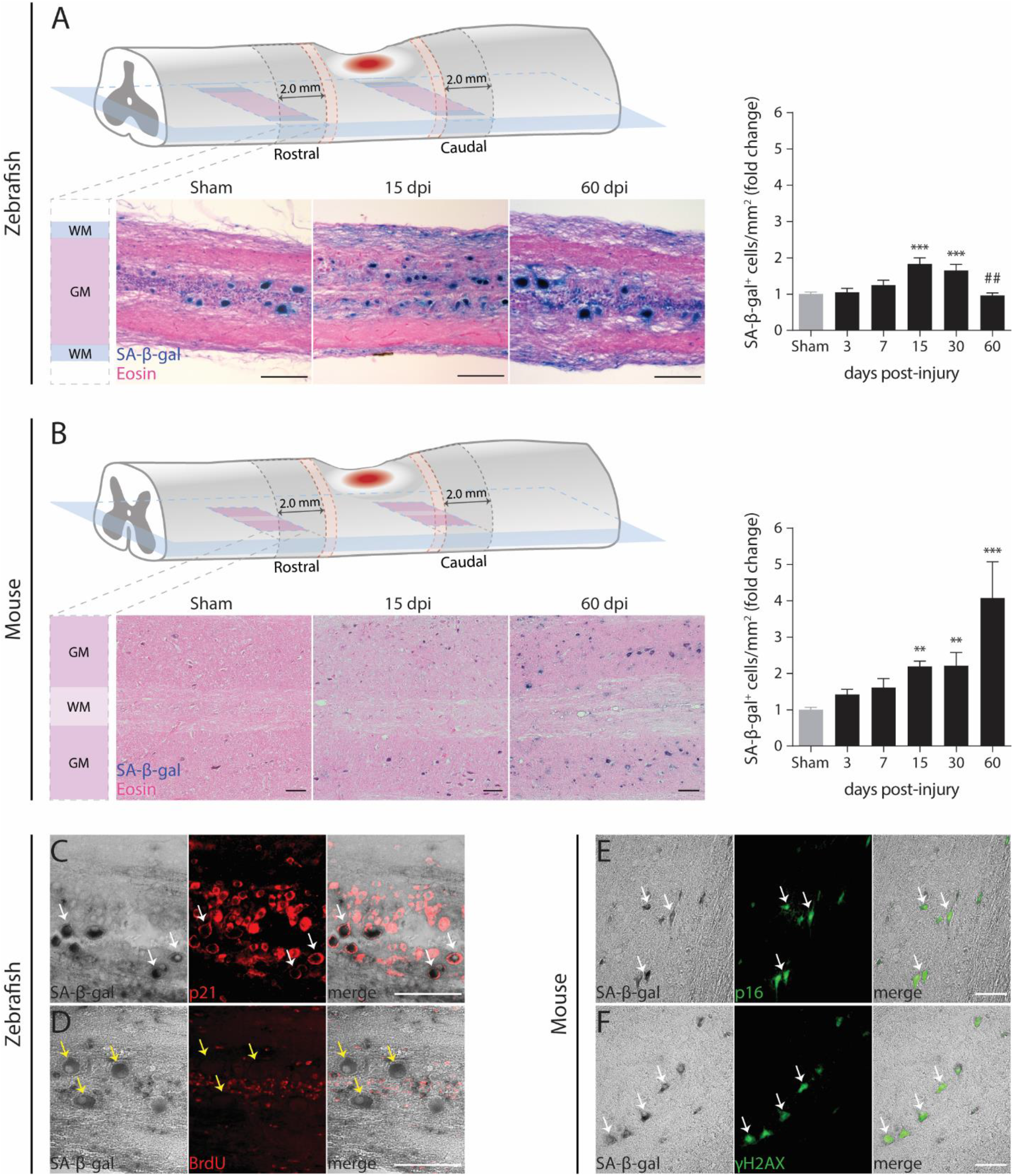
Different senescent cell dynamics are induced after spinal cord injury in zebrafish and in mouse. (**A**) SA-β-gal^+^ cells (blue) were detected and quantified in non-injured (Sham) and injured zebrafish spinal cords at different time-points (3, 7, 15, 30 and 60 days-post-injury, dpi). *n* = 5-16. (**B**) Similarly, SA-β-gal^+^ cells were detected and quantified in mouse laminectomized (Sham) and injured spinal cords at the same time-points. *n* = 4-16. Eosin counterstaining was performed after cryosectioning. Cells were quantified at the lesion periphery along 2.0 mm in longitudinal sections. A 0.5 mm interval (red dashed zone) was established between the lesion border and the beginning of the quantification region. SA-β-gal^+^ cells were only quantified in the gray matter (GM) and not in the white matter (WM). Quantifications are presented in fold change towards Sham. In zebrafish, SA-β-gal^+^ cells reach a peak at 15 dpi (238.6 cells/mm^2^), a two-fold increase compared to Sham (127.2 cells/mm^2^). In mouse, SA-β-gal^+^ cells display a four-fold increase at 60 dpi (80.0 cells/mm^2^) towards Sham (19.0 cells/mm^2^). Data are presented as mean ± SEM. *\*p<0.05, **p<0.01, ***p<0.001*, versus Sham. ^##^*p<0.01,* 60 versus 30 dpi. Scale bars: 100 μm. (**C** and **D**) In zebrafish, SA-β-gal^+^ cells co-localized with the senescence biomarker p21 and were devoid of the proliferation marker BrdU. (**D** and **E**) In mouse a similar co-localization was found between SA-β-gal^+^ cells and the senescence biomarkers p16 and γH2AX. In (**C-E**), images were taken at 15 dpi. White arrows point to representative examples of co-localization. Scale bars: 100 μm.

### SA-β-gal^+^ cells in the zebrafish and mouse spinal cord are mostly neurons

To identify which cell types comprise the SA-β-gal^+^ population, we searched for co-localization with cell-type specific markers. Strikingly, we observed that most SA-β-gal^+^ cells detected in the grey matter at the lesion periphery co-localized with known neuronal markers, namely HuC/D (Figure 2A) and NeuN (Figure 2B). In injured zebrafish spinal cords, 87.8% to 89.1% of SA-β-gal^+^ cells detected in the grey matter co-localized with HuC/D at 15 and 60 dpi, respectively (Figure 2C). Similarly, in mouse spinal cords, 94.8% to 96.0% of SA-β-gal^+^ cells detected in the grey matter co-localized with the pan-neuronal marker NeuN at 15 and 60 dpi, respectively. When we calculated the percentage of SA-β-gal^+^ cells of the total neuronal population at the lesion periphery, we again observed two distinct profiles between these two models. In zebrafish, the percentage of total neurons that are SA-β-gal^+^ reaches a peak of 8.9% at 15 dpi and then returns to basal levels (2.3%) at 60 dpi. On the other hand, mice display 25.3% of senescent neurons at 15 dpi (Figure 2D) and strikingly, this number keeps increasing until 60 dpi, reaching 35.3% of total neurons.

**Fig. 2.**
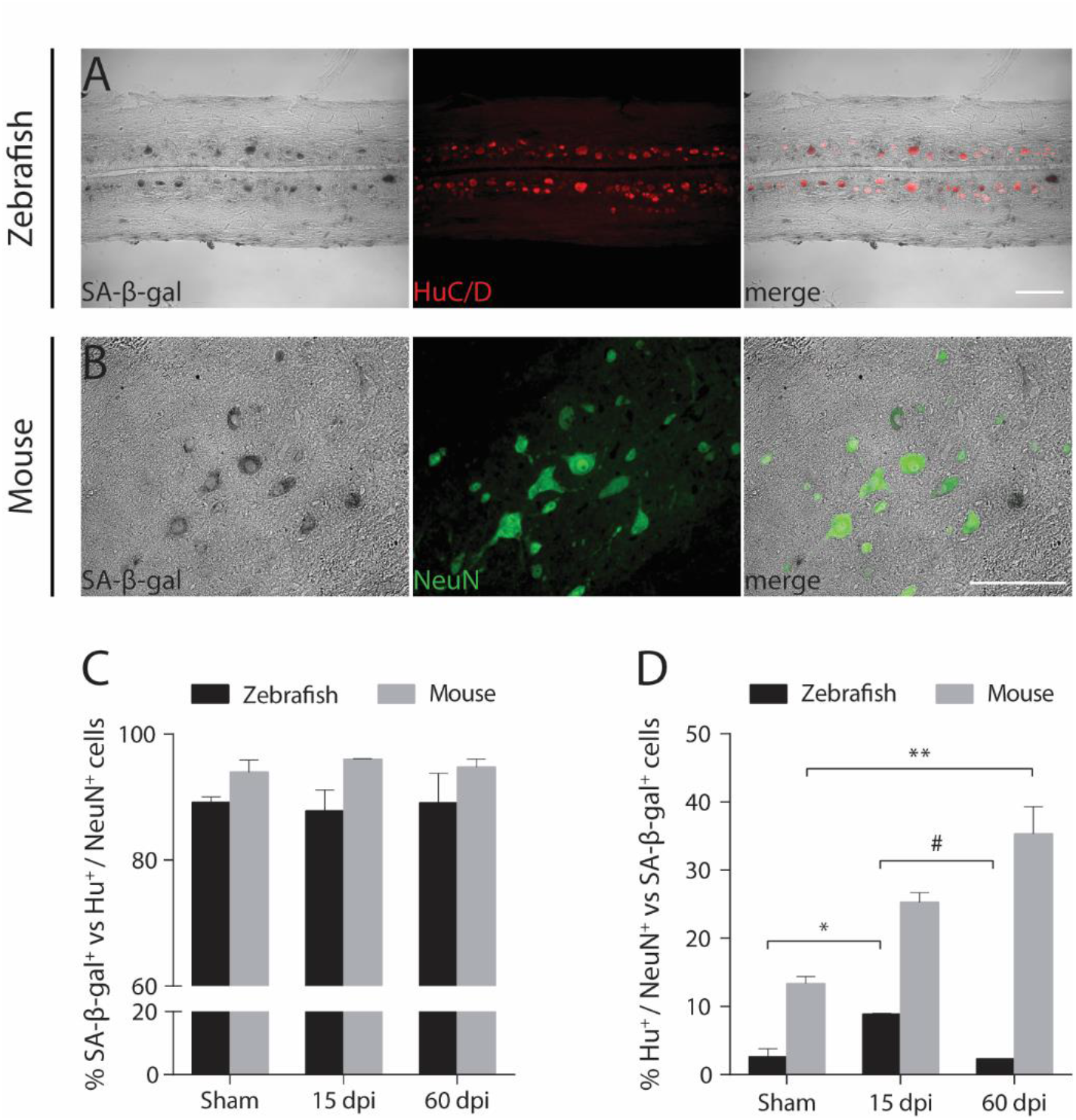
Different profiles of SA-β-gal^+^ neuronal populations between zebrafish and mouse. In zebrafish and mouse, SA-β-gal^+^ cells (black) co-localized with the neuronal markers (**A**) HuC/D (red) and NeuN (green), respectively. Representative images were taken at 15 dpi. In (**C**) and (**D**), the percentages of total SA-β-gal^+^ cells that are HuC/D^+^ or NeuN^+^ and of total HuC/D^+^ or NeuN^+^ neurons that are SA-β-gal^+^ are compared between both models. Cells were quantified at the lesion periphery along 2.0 mm in longitudinal sections. A 0.5 mm interval (red dashed zone) was established between the lesion border and the beginning of the quantification region. Quantifications are presented for uninjured/laminectomized (Sham) zebrafish/mice and at 15 and 60 days post-injury (dpi). *n* = 2-4. Data are presented as mean ± SEM. *\*p<0.05, **p<0.01,* versus Sham. *#p<0.05,* versus 15 dpi. Scale bars: 100 μm.

### Elimination of senescent cells with a senolytic drug improves motor, sensory and bladder functions in a mouse spinal cord contusion injury model

The role of senescence in diverse biological contexts is still poorly understood. Yet, it is accepted that accumulation or persistence of SCs and subsequent chronic exposure to their senescence-associated secretory phenotype (SASP) contribute to loss of tissue function and diminish the repair capacity in aged tissues (Acosta *et al.*, 2013; Campisi, 2013; Childs *et al.*, 2015; Calcinotto *et al.*, 2019).

We hypothesized that accumulation of SCs in the mouse spinal cord is an important factor of the inhibitory microenvironment that undermines the regenerative potential after an injury. To evaluate the impact of eliminating SCs in a mouse contusion model of spinal cord injury, we used the ABT-263 drug, known to work as a powerful senolytic (Paez-Ribes *et al.*, 2019). We administered ABT-263 within the first 14 dpi (sub-acute injury phase), during which the blood-spinal cord barrier remains leaky (Whetstone *et al.*, 2003), thus ensuring maximum accessibility of this drug to the tissue (Fig. 3A). We confirmed that ABT-263 administration by oral gavage from 5 to 14 dpi reduced the number of SA-β-gal^+^ cells in the spinal cord at the lesion periphery, when compared to vehicle administration (**Fig. S2**), thus confirming its senolytic activity in the mouse spinal cord.

**Fig. 3.**
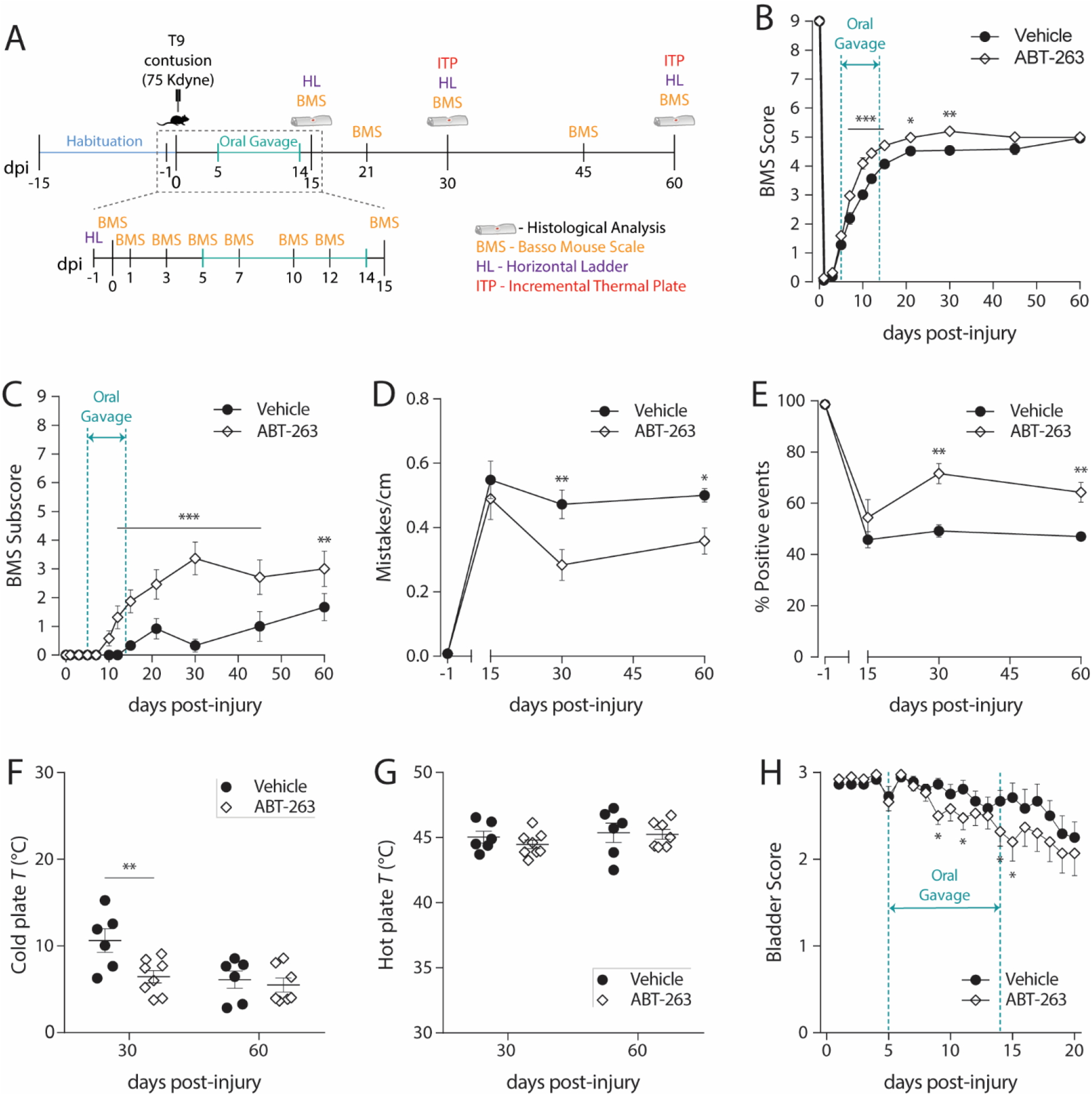
Depletion of senescent cells with ABT-263 improves motor, sensory and bladder function recovery following a spinal cord injury in mice. (**A**) Schematic of the experimental setup. Animals were habituated to the different behavioral setups for a 15-day period, before being submitted to a moderate-to-severe (force: 75 Kdyne; displacement: 550-750 μm) T9 contusion injury. Injured animals received daily vehicle or ABT-263 via oral gavage, from 5 to 14 days-post-injury (dpi). (**B** and **C**) Basso Mouse Locomotor Scale (BMS) score and subscore were evaluated in an open field at different time-points (0, 1, 3, 5, 7, 10, 12, 15, 21, 30, 45 and 60 dpi). *n* = 18-19. (**D** and **E**) The locomotor performance in the Horizontal Ladder (HL) was assessed at −1 (control), 15, 30 and 60 dpi by quantifying the total number of mistakes per centimeter and the percentage of singular positive events (plantar step, toe step and skip) measured and averaged across three successful trials. *n* = 3-6. (**F** and **G**) Thermal allodynia was tested at 30 and 60 dpi by determining the temperature at which injured mice reacted to a cold or hot stimulus. *n* = 6-8. (**H**) Bladder function was grossly evaluated by attributing a bladder score to the amount of urine collected each time a bladder was manually voided. *n* = 18-19. Data are presented as mean ± SEM. *\*p<0.05, **p<0.01, ***p<0.001*, ABT-263 versus Vehicle.

Before injury, all mice presented a normal locomotor behavior in the open field test, which corresponds to the maximum score of 9 in the Basso Mouse Scale (BMS) (Basso *et al.*, 2006). After injury, most injured-mice exhibited complete hindlimb paralysis at 1 dpi and gradually improved locomotor ability reaching a plateau at around 21 dpi (Fig. 3B), similarly to what has been previously described for a contusion injury model in C57BL/6J mice (Basso *et al.*, 2006). In ABT-263 treated animals, BMS scores were significantly higher from 7 until 30 dpi (Fig. 3B) and BMS subscores were significantly higher from 12 to 60 dpi (Fig. 3C), when compared with animals treated with vehicle. Strikingly, at 30 dpi, all ABT-263-treated mice achieved frequent plantar stepping with 93% (14 out of 15) of mice displaying parallel placement of both hindpaws at initial contact and 40% (6 out of 15) also at lift off. Remarkably, 33% (5 out of 15) of mice treated with ABT-263 exhibited consistent plantar stepping and mild trunk stability, one animal showed normal trunk stability with mostly coordinated fore-hindlimb walking, and a second animal displayed some fore-hindlimb coordination, improvements never achieved in vehicle-treated mice.

A finer assessment of locomotion was performed using the Horizontal Ladder (HL) test. Prior to the injury, all mice completed the HL with few to no mistakes or negative events (Fig. 3D, E). Animals treated with ABT-263 made significantly fewer stepping mistakes (Fig. 3D) and displayed significantly more positive stepping events (Fig. 3E) at 30 and 60 dpi, when compared with vehicle-treated mice, thus largely corroborating the results obtained in the open-field.

Thermal allodynia, i.e. hypersensitivity to normally non-noxious stimuli, is a common pain-related symptom associated with spinal cord injuries (Nakae *et al.*, 2011; Watson *et al.*, 2014). Using an incremental thermal plate (ITP), we could compare the temperature threshold necessary to elicit an avoidance behavior to a cold or a hot stimulus between the two experimental groups. Considering that uninjured C57BL/6J mice only exhibit nocifensive reaction to cold between 2 to 4°C (Yalcin *et al.*, 2009), ABT-263 treatment significantly decreased cold hypersensitivity at 30 dpi, compared to vehicle-treated animals who showed an average temperature reaction to cold of 10.6°C (Fig. 3F). We found no effect of ABT-263 administration on the threshold temperature required to prompt a nocifensive reaction to a hot stimulus, when compared to vehicle administration (Fig. 3G).

Another common consequence of spinal cord injury is bladder dysfunction (Yoshimura, 1999). We assessed bladder function by attributing a score to the amount of urine retained each day. In contrast to injured mice treated with vehicle, injured-mice treated with ABT-263 exhibited smaller volumes of retained urine from 9 to 20 dpi (Fig. 3H), an effect that was lost after 20 dpi (data not shown).

In a completely independent study, we eliminated SCs with a cocktail of two different drugs, Dasatinib plus Quercetin (D+Q), known to have a strong senolytic activity (Zhu *et al.*, 2015), using the same injury parameters and administration time-window for the sake of comparison (**Fig. S3A-C**). D+Q-treated animals exhibited a significantly improved locomotor function after a spinal cord injury towards vehicle-treated mice, resulting in higher BMS scores (**Fig. S3D, E**) and improved HL performances (**Fig. S3F, G**). Strikingly, similarly to ABT-263, D+Q administration also decreased the hypersensitivity to a cold stimulus at 30 dpi (**Fig. S3H**), while no effects were observed in response to a hot stimulus (**Fig. S3I**). These results corroborate those obtained with ABT-263, reinforcing the positive effect of eliminating SCs on locomotor and sensory recovery after a spinal cord injury in mammals. In accordance, the D+Q cocktail significantly decreased the number of SA-β-gal^+^ cells at the lesion periphery (**Fig. S3J**), thus confirming its senolytic effect in the mouse spinal cord.

### The senolytic ABT-263 promotes myelin preservation after spinal cord injury

After a spinal cord injury, oligodendrocytes undergo both necrotic and apoptotic cell death, which results in demyelination around the lesion, impairing function of unprotected fibers, contributing to the accumulation of cell debris and potentiating the inhibitory microenvironment for repair (Crowe *et al.*, 1997; Emery *et al.*, 1998; Totoiu and Keirstead, 2005). To evaluate the effect of eliminating SCs on the demyelination status after injury, we used the FluoroMyelin^TM^ Green fluorescent myelin staining to compare the spared white matter area per total cross-sectional area (% CSA) between ABT-263-treated mice and vehicle-treated mice along 2 mm around the lesion epicenter. Remarkably, ABT-263 treatment consistently resulted in significantly greater white matter sparing levels across all experimental time-points, an effect that at 30 dpi was more prominent at the caudal side of the lesion (Fig. 4A-C).

**Fig. 4.**
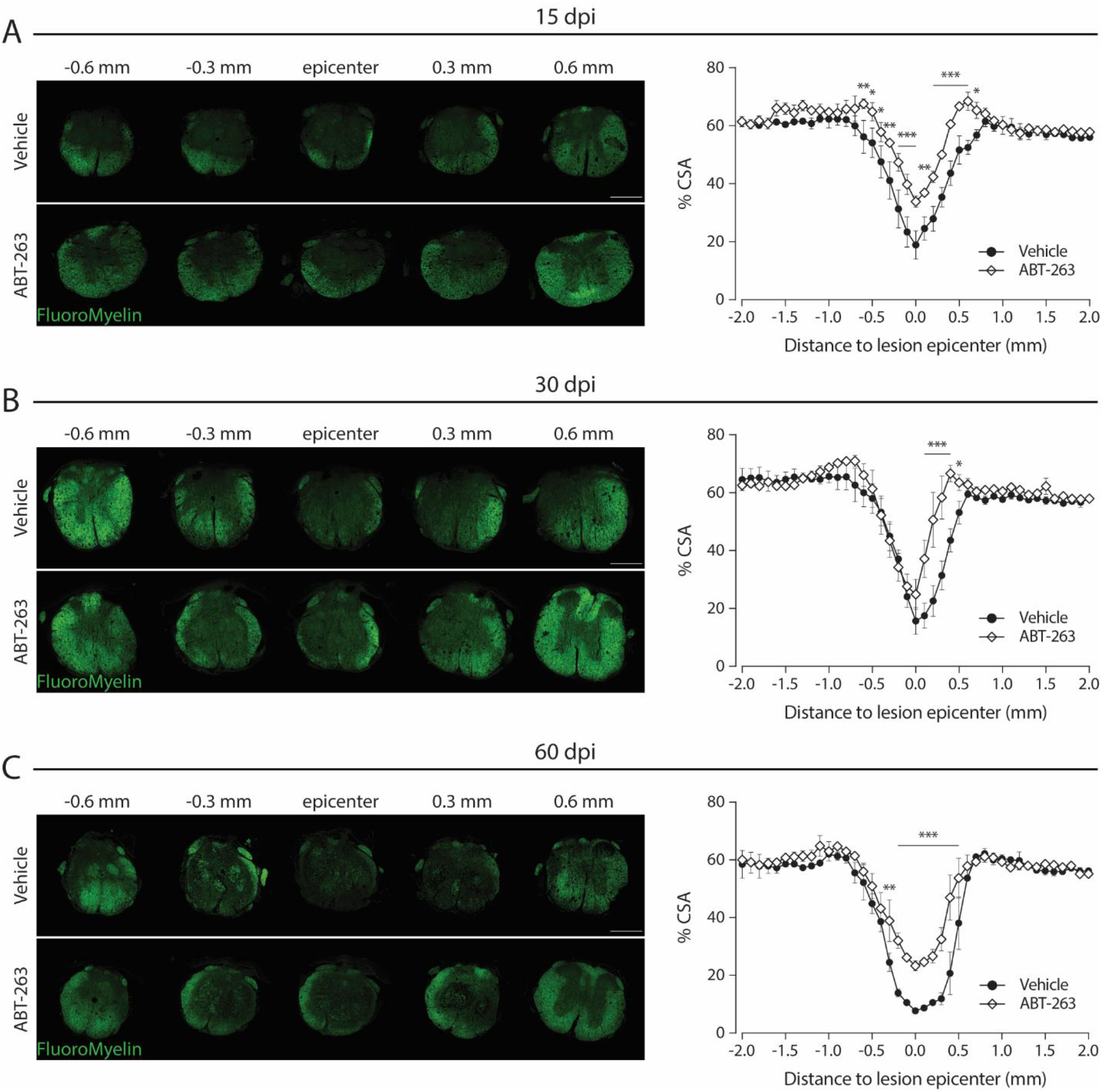
White matter sparing is increased after eliminating senescent cells with ABT-263. Transversal sections at different distances from the lesion epicenter of an injured spinal cord (**A**) at 15 days-post-injury (dpi), (**B**) at 30 dpi and (**C**) at 60 dpi, treated with Vehicle and ABT-263 and stained with FluoroMyelin (green) and the corresponding quantifications. In (**A-C**), white matter sparing was assessed by normalizing the area stained with FluoroMyelin (green) to the total cross-sectional area (CSA) of spinal cord sections every 100 μm ranging from 2 mm rostral and 2 mm caudal to the lesion epicenter. Scale: 500 μm. *n* (15 dpi) = 3-4; *n* (30 dpi) = 3-4; *n* (60 dpi) = 2-3. Data are expressed as % CSA and presented as mean ± SEM. *\*p<0.05, **p<0.01, ***p<0.001*, ABT-263 versus Vehicle.

### ABT-263 supports axonal growth after spinal cord injury

The normal neural circuit organization is disrupted after a spinal cord injury. However, spared neural tissue can, to a certain extent (depending on the severity of the lesion), reorganize itself in order to establish new lines of communication across and beyond the injury (Courtine and Sofroniew, 2019). This plasticity potential explains why after an incomplete lesion (e.g. our contusion injury model) mice can partially restore their locomotor function.

We hypothesized that eliminating SCs would provide a more favorable environment for axonal preservation and growth after a spinal cord injury. To assess this, we tested the expression of neuronal growth-associated protein 43 (GAP43), which is highly expressed in neuronal growth cones during development and axonal regeneration (Benowitz and Routtenberg, 1987). We performed immunostainings for GAP43 in dorsal-ventral longitudinal sections spanning the ventral horn (where the motor neurons are located) and quantified the number of GAP43^+^ fibers at specific distances from the lesion epicenter (Fig. 5A), as previously described (Hata *et al.*, 2006; Almutiri *et al.*, 2018). At 30 dpi, ABT-263-treated mice had a significant increased number of GAP43^+^ axons across most inspected regions, including at the lesion epicenter, compared to vehicle-treated mice (Fig. 5B, C).

**Fig. 5.**
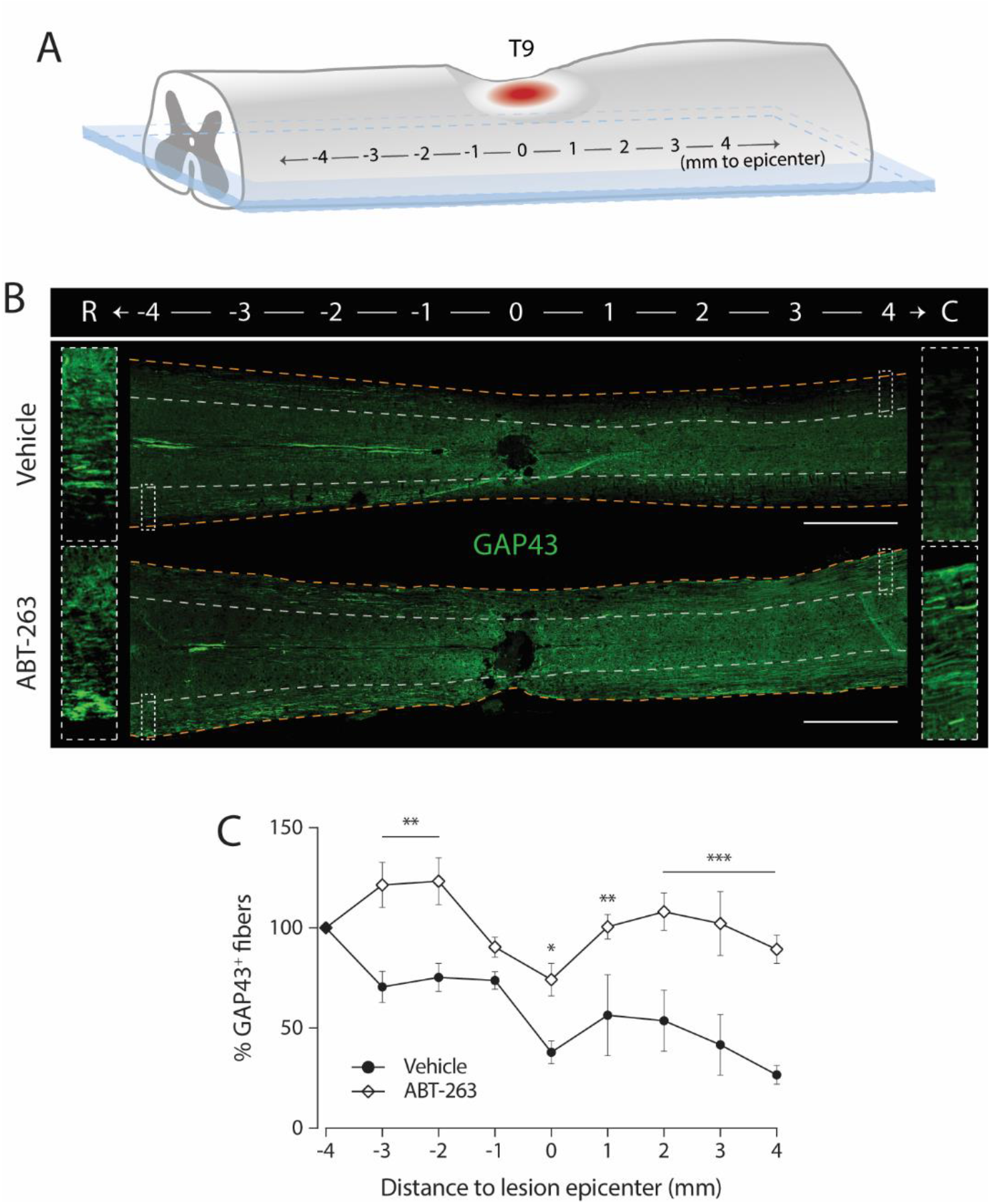
ABT-263 administration increases the number of GAP43^+^ fibers. (**A**) Longitudinal coronal sections of the ventral region of the spinal cord (delineated in blue) were obtained. GAP43^+^ axonal fibers were counted at specific distances (every 1 mm) from the lesion epicenter along the rostral (R)-caudal (C) axis of injured spinal cords. (**B**) Longitudinal coronal section of the ventral region of a spinal cord treated with Vehicle or ABT-263 and stained with GAP43. GAP43^+^ axonal fibers were counted at specific distances (every 1 mm) from the lesion epicenter along the rostral (R)-caudal (C) axis. Fibers were quantified only in the white matter in the ventral region of the spinal cord (delineated in blue). Light grey dashed lines delimitate the white matter from the grey matter. Orange dashed lines outline the limit of the spinal cord section. Scale: 1000 μm. (**C**) Axon number was calculated at 30 days post-injury (dpi) as a ratio of the total GAP43^+^ fibers at 4 mm rostral (−4) from the lesion epicentre (0). *n* = 3. Data are presented as mean ± SEM. \**p*<0.05, \*\**p*<0.01, \*\*\**p*<0.001, ABT-263 versus Vehicle.

### Administration of the senolytic ABT-263 reduces the fibrotic scar

Scar formation following spinal cord injury constitutes a major barrier for axonal regrowth (Jared M Cregg *et al.*, 2014). Inside the lesion core, a subset of proliferating PDGFRβ^+^ perivascular cells give rise to a fibrotic scar with dense deposition of extracellular matrix components (Soderblom *et al.*, 2013). In fact, it has been shown that reducing the pericyte-derived fibrotic scar facilitates functional recovery after spinal cord injury in mice (Dias *et al.*, 2018). We examined the size and length of the fibrotic scar using a double immunostaining with PDGFRβ (a pericyte marker that labels the fibrotic component of the scar) and GFAP (a glial marker that helps delineate the fibrotic scar compartment). At the lesion epicenter, mice treated with the senolytic exhibited a significantly reduced PDGFRβ^+^ area at 15 and 60 dpi when compared to mice treated with vehicle (Fig. 6A-D). Using the same double immunostaining, we were able to define the extension of the scar by tracing, rostrally and caudally to the epicenter, signs of fibrotic PDGFRβ^+^ staining in the dorsal side of the spinal cord (**Fig. S4A**). With this analysis, we observed that the total length of the fibrotic scar was shorter at 15 dpi in ABT-263-treated mice, an effect sustained until 30 dpi only at the caudal side (**Fig. S4B**).

**Fig. 6.**
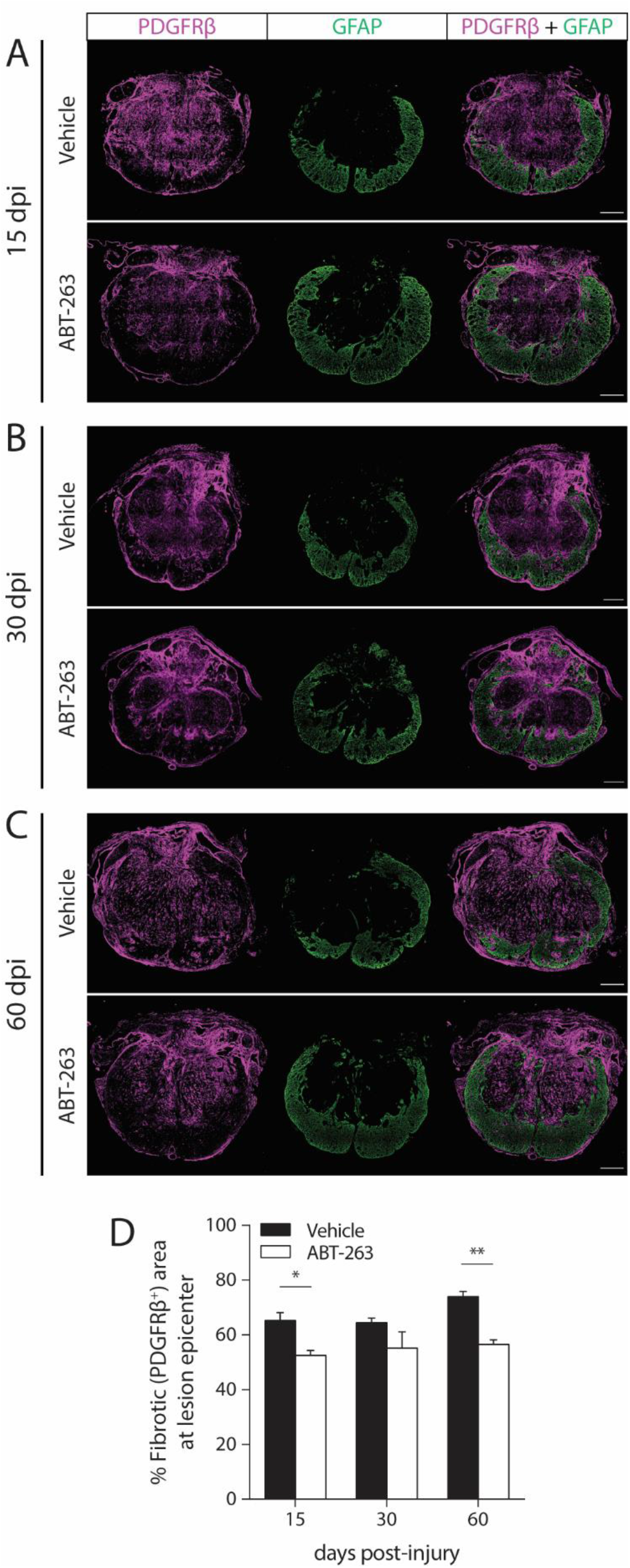
Eliminating senescent cells leads to a reduction of the fibrotic scar. Transversal sections at the lesion epicenter of an injured spinal cord (**A**) at 15 days-post-injury (dpi), (**B**) at 30 dpi and (**C**) at 60 dpi treated with Vehicle and ABT-263 and stained with the fibrotic scar marker PDGFRβ^+^ area (magenta) and with the astrocytic scar marker GFAP (green). The fibrotic scar area was evaluated by normalizing the PDGFRβ^+^ area (magenta) to the total cross-sectional area at the lesion epicenter. GFAP^+^ tissue (green) surrounds the fibrotic core. Scale: 200 μm. *n* (15 dpi) = 3-4; *n* (30 dpi) = 3-4; *n* (60 dpi) = 2-3. (**D**) The percentage of fibrotic tissue in the injury core was quantified at 15, 30 and 60 dpi. Data are presented as mean ± SEM. \**p*<0.05, \*\**p*<0.01, ABT-263 versus Vehicle.

### Macrophage numbers at the injury site are reduced following ABT-263 treatment

A spinal cord lesion in mice elicits a strong and long-lasting inflammatory response that potentiates secondary injury (Blight, 1985; Popovich, Wei and Stokes, 1997). Macrophages are the most abundant inflammatory cells in a spinal lesion, infiltrating the injury core and releasing several molecules, namely nitrogen/oxygen metabolites, cytokines, proteases and chondroitin sulphate proteoglycans that can cause cellular damage and inhibit axonal growth (Fitch and Silver, 1997). Importantly, depletion of macrophages was demonstrated to promote repair and partial motor recovery after spinal cord injury in rats (Popovich *et al.*, 1999). Additionally, senescence is closely linked to inflammation. SCs, through their SASP, can secrete a plethora of immune modulators and proinflammatory cytokines like TNF-α and CCL2 (two potent macrophage recruiters), interleukin-6 (IL-6), IL-8 and IL-1α (Coppé *et al.*, 2010). Therefore, the accumulation/persistence of SCs in tissues is usually associated with chronic inflammation. To investigate the impact of the accumulation of SCs on inflammation in the mouse spinal cord after an injury, we performed immunostainings with the pan-macrophage marker F4/80. As anticipated, by eliminating SCs with ABT-263, we observed lower levels of inflammatory macrophages (% F4/80^+^ area) in spinal cord sections spanning the lesion area, particularly at 30 dpi (Fig 7A-C).

**Fig. 7.**
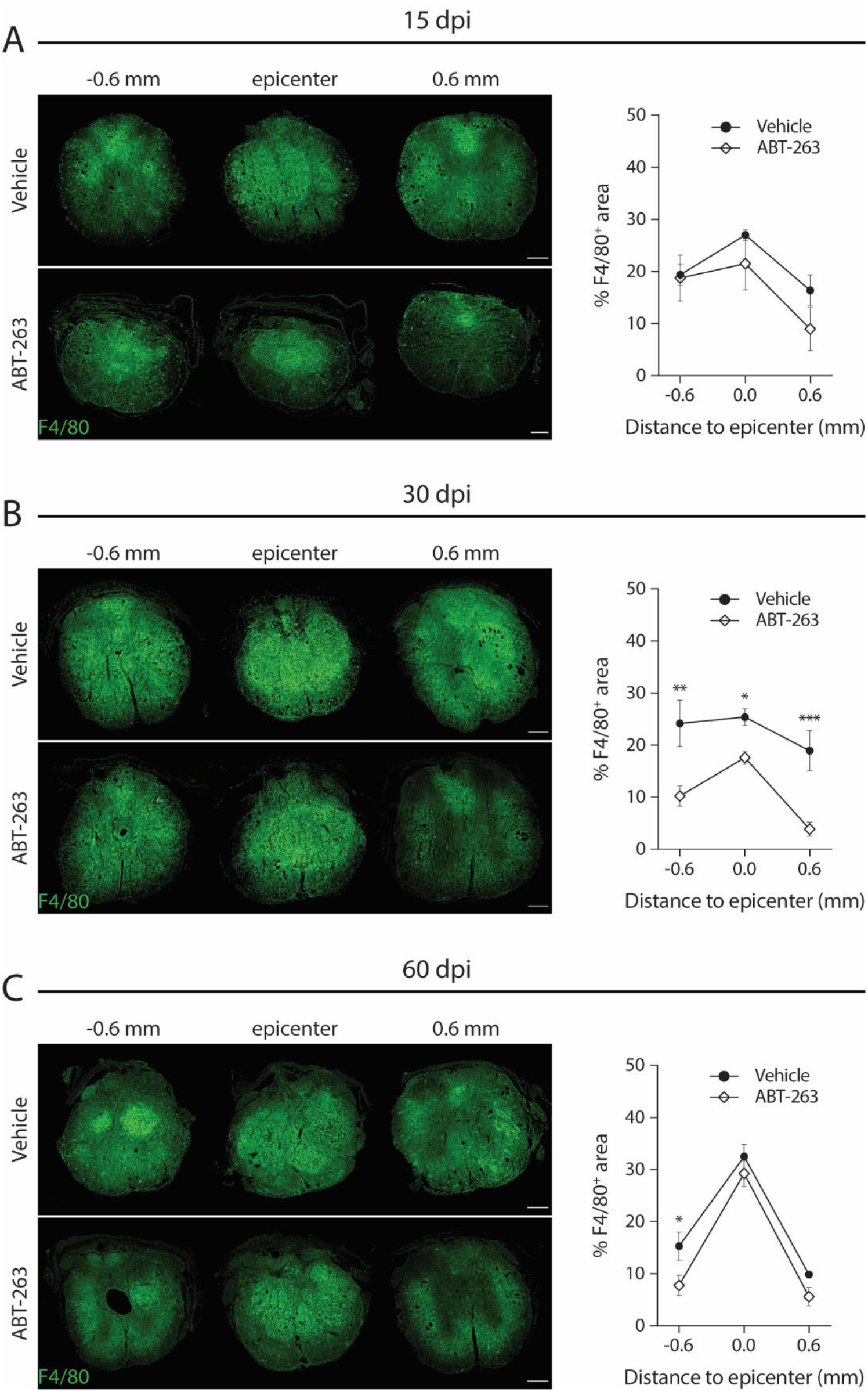
ABT-263 treatment decreases the number of macrophages at the injury site. Transversal sections at the lesion epicenter of an injured spinal cord (**A**) at 15 days-post-injury (dpi), (**B**) at 30 dpi and (**C**) at 60 dpi treated with Vehicle and ABT-263 and stained with the pan-macrophage marker F4/80 (green). In (**A-C**), the area of F4/80^+^ tissue was measured at lesion epicenter and 600 μm rostral and caudal from the epicenter. Measurements are expressed as a percentage of the total cross-sectional area. *n* (15 dpi) = 3-4; *n* (30 dpi) = 3-4; *n* (60 dpi) = 2-3. Scale: 200 μm. Data are presented as mean ± SEM. \**p*<0.05, \*\**p*<0.01, \*\*\**p*<0.001, ABT-263 versus Vehicle.

## DISCUSSION

The concepts of cellular senescence and their SASP have evolved remarkably over the last years. Yet, most of the mechanisms underlying the complexity of each senescent program remain unknown. While further studies are necessary to understand and reconcile the physiological and pathological roles of senescence, current knowledge favors the premise that transient and controlled induction of SCs is beneficial whereas accumulation and persistence of SCs is detrimental (Rhinn, Ritschka and Keyes, 2019). Interestingly, the role of senescence in wound repair and organ regeneration contexts is still quite uncharted ground.

Here, we describe the induction of SCs as a new cellular response triggered by an injury in the spinal cord. Our data shows that the majority of senescent cells, quantified in the grey matter located at the lesion periphery of injured spinal cords, are neurons. Post-mitotic neurons exhibiting several senescence features have already been described in the rat cortex in both rodent and human aging brains (Jurk *et al.*, 2012; Chinta *et al.*, 2015; Kang *et al.*, 2015; Moreno-Blas *et al.*, 2019; Walton and Andersen, 2019). Moreover, a recent study has demonstrated that senescent cortical neurons have their own SASP that is able to induce paracrine senescence in mouse embryonic fibroblasts (Moreno-Blas *et al.*, 2019).

In the regenerating zebrafish model, SCs start to accumulate essentially at the lesion periphery but are eventually cleared and returned to basal levels. This transient profile of SCs induction seems to be a conserved injury-response in organs with regenerative capacities, since it was also described in amputated appendages and damaged hearts of zebrafish, salamanders and neonatal mice (Yun, Davaapil and Brockes, 2015; Da Silva-Álvarez *et al.*, 2019; Sarig *et al.*, 2019). In the non-regenerating mouse however, the induced SCs persist at the lesion periphery and do not show any signs of being efficiently reduced over time. How are these transient vs. persistent senescent profiles established is not known, but it is possible that these are associated with a specific SASP with different capacities to support a cell clearance mechanism. This seems to be the case in the salamander regenerative limb paradigm, where macrophages were shown to be an essential part of the mechanism that eliminates SCs (Yun, Davaapil and Brockes, 2015). The functional demonstration of the positive role of transient SCs in a regenerative context comes from the observation that reducing SCs leads to a regeneration delay of amputated pectoral fins in zebrafish (Da Silva-Álvarez *et al.*, 2019). These findings are in line with what was previously reported for skin wound healing, where transient SCs were found to be fundamental for tissue remodeling and repair (Demaria *et al.*, 2014).

Even though there is currently no tool to efficiently eliminate SCs in the zebrafish spinal cord, the observed transient profile suggests that a timely clearance of SCs might be required for proper regeneration to occur. In line with this idea, the persistent senescence profile we found in mice is compatible with the inability of mammals to regenerate the spinal cord after a lesion. Consequently, investigating the functional role of induced-SCs after a spinal cord injury in mice becomes of paramount importance, as it could create new opportunities for therapeutic interventions.

Senolytic drugs selectively eliminate SCs by transiently disabling the pro-survival networks. One of such drugs is ABT-263, a specific inhibitor of anti-apoptotic proteins BCL-2 and BCL-xL, already shown to selectively and efficiently kill senescent cells *in vivo* in mice (Demaria *et al.*, 2014; Chang *et al.*, 2016). We used ABT-263 to eliminate SCs after a spinal cord T9 contusion injury in mouse and its effects on motor, sensory and bladder function recovery were evaluated. We targeted the elimination of SCs in the subacute injury phase to guarantee that we were acting when macrophage infiltration, reactive astrogliosis and scar formation are taking place (Siddiqui, Khazaei and Fehlings, 2015) but also to prevent their accumulation at the lesion periphery which becomes statistically significant already at 15 dpi. We were able to show that SCs were eliminated in injured mice treated with ABT-263, with elimination efficiencies similar to what was previously described (Demaria *et al.*, 2014). Although the number of SCs are indeed reduced in all time-points analyzed, they start to slowly re-emerge after the end of ABT-263 administration. This might be a consequence of paracrine senescence, a SASP-mediated event where the remaining SCs can induce senescence in nearby cells (Herranz and Gil, 2018). In fact, as a Bcl-2/BCL-xL inhibitor, ABT-263 induces apoptosis in existent SCs, but does not prevent the induction of new SCs. This becomes relevant when thinking in a translational approach where the administration time window should be carefully established for high-efficacy with low-toxicity.

The treatment of spinal cord injured mice with the senolytic ABT-263 significantly improved locomotor performance in BMS and HL tests, an effect maintained until the end of the study (i.e. 60 dpi), and also bladder function during the administration period. Interestingly, at 30 dpi, ABT-263-treated animals showed a normal sensitivity to a non-noxious cold stimulus but no effects were observed upon a hot stimulus. This may indicate that SCs and their SASP are acting through specific neural substrates, namely through transient receptor potential member 8 (Trpm8) cation channels-primary molecular transducers of cold somatosensation (Ran, Hoon and Chen, 2016).

Importantly, the effects of ABT-263 on locomotor and sensory recovery were corroborated by a second independent assay using the Dasatinib plus Quercetin (D+Q) senolytic cocktail, know to exert its activity preferentially via PI3K inhibition Zhu *et al.*, 2015). In addition, we could also show that, similarly to ABT-263, the D+Q cocktail also resulted in an efficient depletion of SCs at the spinal cord lesion periphery. Together, these results highlight the detrimental impact of the persistent accumulation of SCs on motor and sensory functions after a spinal cord injury.

Persistent senescent fibroblasts and myogenic cells through their SASP were shown to promote a pro-fibrotic response and to limit tissue repair in fibrotic lung disease (Schafer *et al.*, 2017) and injured muscles (Le Roux *et al.*, 2015), respectively. Accordingly, we showed that the effect of ABT-263 on SCs depletion was translated into a consistently reduced fibrotic scar area and length. In addition, SCs depletion with ABT-263 results in a higher myelin preservation over time. While decreasing demyelination helps preserve the function of spared axons, a smaller scar provides a better microenvironment for the reorganization of spared axons around the lesion (Courtine and Sofroniew, 2019). Consistent with this scenario, ABT-263 treatment promoted an increased expression of the growth-associated GAP43 protein. Altogether, these effects are likely underlying the locomotor improvements observed. Noteworthy, the effects of ABT-263 seems to be more pronounced at the caudal side of the lesion. These differences in lesion responses to treatment suggest the existence of different SASP programs between the rostral and caudal sides, something that requires further investigation in the future.

The neuroinflammatory response after a spinal cord injury worsens throughout the secondary damage phase, becomes chronic and is associated with neurotoxicity (Fleming *et al.*, 2006). Preventing the accumulation of SCs during the subacute injury phase with the administration of ABT-263, led to a reduction in the number of inflammatory macrophages and concomitantly to a better functional outcome in injured mice. Interestingly, persistent SCs are known to create a chronic inflammatory tissue microenvironment by secreting proinflammatory cytokines like IL-1α, IL-1β, IL-6, IL-8 or TNF-α, which are all well-established components of the SASP (Coppé *et al.*, 2008, 2010). Moreover, neutralization of IL-1β and TNF-α signaling has already been shown to improve functional recovery after spinal cord injury (Nesic *et al.*, 2001; Genovese T, Mazzon E, Crisafulli C, Di Paola R, Muià C, Esposito E, Bramanti P, 2008).

Our data provides evidence for the remarkable beneficial outcomes of eliminating SCs in the context of a spinal cord injury, namely by reducing inflammation, limiting scaring, preserving myelin and allowing axonal growth. Targeted elimination of SCs emerges as a promising therapeutic approach to promote functional repair of an injured spinal cord, repurposing the use of senolytic therapies already under clinical trials for cancer and age-related disorders (Paez-Ribes *et al.*, 2019). Given the extreme complexity and multifaceted mechanisms underlying spinal cord repair, it is doubtful that, by itself, a senolytic drug can generate clinically meaningful functional improvements. We do believe though that this strategy shows great potential to be combined with other existing biological and engineering approaches in a combinatorial therapeutic logic.

## MATERIALS AND METHODS

### Ethics statement

All handling, surgical and post-operative care procedures were approved by Instituto de Medicina Molecular Internal Committee (ORBEA) and the Portuguese Animal Ethics Committee (DGAV), in accordance with the European Community guidelines (Directive 2010/63/EU) and the Portuguese law on animal care (DL 113/2013). All efforts were made to minimize the number of animals used and to decrease suffering of the animals used in the study.

### Study Design

#### Rationale and experimental design

This study was designed to investigate the role of senescence in a spinal cord injury context. Standard senescence biomarkers (SA-β-gal, p21^CIP1^, p16^INK4a^ and γH2AX) were used to characterize SCs induced after injury. We used ABT-263, a drug with powerful senolytic activity, to pharmacologically deplete SCs during the sub-acute phase of the injury. BMS and HL tests were used to study locomotor recovery, while the ITP assessed sensory function. At the cellular level, we evaluated the impact of eliminating SCs on myelin and axonal preservation, fibrosis and inflammation.

#### Animals

AB strain zebrafish (*Danio rerio*) were obtained from Zebrafish International Resource Center (ZIRC). Animals were bred, grown and maintained on a 14-hour/10-hour light/dark cycle at 28°C following the standard guidelines for fish care and maintenance protocols. Male and female fish were used in the experiments.

Adult (8-9 weeks old) female C57BL/6J mice (*Mus musculus*) were purchased from Charles River Laboratory. Animals were housed in the Instituto de Medicina Molecular animal facility under conventional conditions on a 12-hour light-dark cycle with *ad libitum* access to food and water.

#### Randomization and blinding

After the injury, animals were randomly assigned to each experimental group and end-point. Experimenters were blinded for the whole duration of the study and data analysis.

#### Sample size and inclusion criteria

Our inclusion criteria depended on our biomechanical and behavioral injury parameters (displacement: 550-750 μm; BMS score ≤ 0.5 averaged across both hindlimbs at 1 dpi). According to these criteria, a total of 19 and 18 mice were used for the ABT-263-treated and vehicle-treated experimental groups, respectively.

#### Selection of endpoints

The selection of endpoints was based on previous studies and pilot experiments in which we characterized both models. We took in account the different phases of the injury progression (subacute and chronic) in a mouse contusion model and the whole regeneration period (60 dpi) of the zebrafish.

### Spinal cord injury (SCI) and post-operative care

#### Zebrafish

Adult zebrafish (6 months old) were anaesthetized in 0.016% tricaine (Sigma, MS222), and a spinal cord crush injury was performed according to a previously described method (Fang *et al.*, 2012). Animals were allowed to recover at 28°C in individual tanks until different experimental time-points (3, 7, 15, 30 and 60 days post-injury, dpi), upon which they were sacrificed and the spinal cords (5-6 mm width) dissected. In control fish, a Sham injury was performed by making an incision at the side of the animal but leaving the spinal cord intact before sealing the wound.

#### Mouse

Before being assigned to SCI, mice went through a two weeks-period of handling and acclimatization, during which body weight was assessed to ensure ideal surgical weight (18-20 g). Animals (10-11 weeks old) were anesthetized using a cocktail of ketamine (120 mg/kg) and xylazine (16 mg/kg) administered by intraperitoneal injection (IPi). For spinal contusion injuries, a laminectomy of the ninth thoracic vertebra (T9), identified based on anatomical landmarks, was first performed (Harrison *et al.*, 2013) followed by a moderate-to-severe (force: 75 Kdyne; displacement: 550-750 μm) contusion using the Infinite Horizon Impactor (Precision Systems and Instrumentation, LLC.) (Scheff *et al.*, 2003). The mean applied force and tissue displacement for each experimental group are shown in **Supplementary Fig. S5A-B**. There were no differences in injury parameters between experimental groups. After SCI, the muscle and skin were closed with 4.0 polyglycolic absorbable sutures (Safil, G1048213). In control uninjured mice (Sham), the wound was closed and sutured after the T9 laminectomy and the spinal cord was not touched. Animals were injected with saline (0.5 ml) subcutaneously, then placed into warmed cages (35°C) until they recovered from anaesthesia and for the following recovery period (3 days). To prevent dehydration mice were supplemented daily with saline (0.5 ml, subcutaneously) for the first 5 dpi. Bladders were manually voided twice daily for the duration of experiments. Body weight was monitored weekly.

### Drug treatment

ABT-263 (Selleckchem, S1001, 50 mg/kg/day) or vehicle (Corn oil, Sigma, C8267) were administered by oral gavage, as described previously (Chang *et al.*, 2016), for 10 consecutive days starting at 5 dpi until 14 dpi. After spinal cord injuries, mice were randomly assigned to each group for each endpoint group, 15 dpi (n = 10), 30 dpi (n = 14) and 60 dpi (n = 13). Within the same cage animals received different treatments to exclude specific environmental cage input. Experimenters were blinded for the whole duration of the study by coding the treatment.

### Behavior assessment

#### Basso Mouse Scale (BMS)

Two weeks before the beginning of the study mice were habituated to the open-field arena to decrease anxiety and distress. On the day of the behavioral test two investigators, blind to treatment, assessed mouse hind limb function and locomotion using the BMS (Basso *et al.*, 2006). Locomotor behavior (BMS scores and subscores) was assessed at 0 (baseline), 1, 3, 5, 7, 10, 12, 15, 21, 30, 45 and 60 dpi.

#### Horizontal Ladder (HL)

On the previous week before SCI, mice were trained to walk along a HL as previously described (Cummings *et al.*, 2007). This task requires mice to walk across a HL that consists of a 60 cm length x 8 cm width transparent corridor with rungs spaced 1 cm apart. A mirror was placed underneath the ladder and mice were video-recorded from the side view to be able to see paw placement on the rung in the mirror. Home cage bedding and/or treat-pellets were placed at the end to stimulate motivation. Ideally, each mouse was able to perform at least three successful trials along the ladder. The three best attempts were scored. A paw falling below the rungs of the ladder during a step in the forward direction was counted as one mistake. The total number of mistakes was averaged across the three trials per mouse and quantified as mistakes per centimetre. The total number of positive and negative events for each rung in each attempt were also quantified and are divided as singular positive events (plantar step, toe step and skip) or singular negative events (slip, miss and drag). Baseline data were assessed 1 day before SCI. Mice were tested on the HL at 15, 30 and 60 dpi.

#### Incremental Thermal Plate (ITP)

Each mouse was placed into the observation chamber of the IITC’s Incremental Hot Cold Plate (IITC Inc. Life Science) with a starting temperature of 37°C, as previously described (Yalcin *et al.*, 2009). The plate was then either heated up to 49°C or cooled down to 0°C at a rate of 6°C per minute until the animal showed nocifensive behavior involving either hindpaw. The typical response was hindpaw licking, shaking and lifting of the paw, jumping and extensor spasm. The plate temperature evoking any of these nocifensive reactions confined to any hindpaw was regarded as the noxious heat/cold threshold of the animal. Following the recording of the threshold temperature, the animal was immediately removed from the plate. The threshold measurement was repeated after 30 minutes and the mean of the two thresholds was considered as the control noxious heat/cold threshold of the animal. Animals were tested in the ITP at 30 and 60 dpi.

#### Bladder function

Bladder function was grossly evaluated by attributing an averaged bladder score (from 0 to 3) to the two daily urine collections, depending on the amount of retained urine (0 – empty bladder; 1 – small bladder; 2 – medium bladder; 3 – large/full bladder). Bladder voiding times, as well as voiding-responsible experimenters, were maintained consistent throughout the experiment.

### Tissue processing

#### Zebrafish

The vertebral column of adult zebrafish was dissected and fixed in 4% paraformaldehyde (PFA) at 4°C overnight (ON). After fixation, the spinal cord was isolated from the vertebral column. Samples were washed 3 times in phosphate-buffered saline (PBS) during the day and incubated ON with SA-β-gal staining solution (see details below). Following the SA-β-gal staining protocol, samples were cryoprotected in 30% sucrose/0.12 M phosphate buffer (PB) for a minimum of 72 hours at 4°C or until the tissue sinks to the bottom of the vial, followed by another embedding in 7.5% gelatin (Sigma, G6144)/15% sucrose/0.12 M PB and subsequently frozen. The samples were cryosectioned in 12 μm-thick longitudinal slices using a Cryostat Leica CM 3050S and either processed for immunohistochemistry or counterstained with eosin for SA-β-gal quantifications.

#### Mouse

Mice were anesthetized with ketamine/xylazine mix (120 mg/kg + 16 mg/kg, IPi) and then transcardially perfused with 0.9% sodium chloride followed by 4% PFA. Post-mortem anatomical assessment of the T9 was confirmed to ensure correct thoracic contusion. Spinal cords were removed, post-fixed in 4% PFA for 2 hours and then incubated ON with SA-β-gal staining solution (see details below). Samples (1 cm in length) were then submitted to the same cryoprotection/embedding and cryosection/staining procedures as for zebrafish spinal cords. Tissue sections were cut in series either transversally (10 μm thick, 10 slides per series) or longitudinally (10 μm thick, 6 slides per series). For each time-point, samples were distributed as equally as possible in cuts along the coronal (rostral-caudal) axis and horizontal (dorsal-ventral) axis. Slides were stored at −0°C until needed. Every block, as well as every slide, was coded until the end of each analysis.

### SA-β-gal staining

SA-β-gal activity was determined in isolated spinal cords using the SA-β-gal kit (Cell Signalling, #9860) according to manufacturer’s instructions, with minor adaptations. Spinal cords were fixed ON in 4% PFA, washed three times in PBS and stained ON at 37°C using the SA-β-gal staining solution (pH 5.9-6.1, prepared according to kit’s instructions). The samples were then washed in PBS, fixed in 4% PFA for 4 hours, washed 3 x 5 minutes in PBS and embedded in sucrose as described above.

### Immunohistochemistry

To perform immunostaining in sections, the gelatin was removed from the cryosections using PBS heated to 37°C (4 × 5-minute washes). After incubation with blocking solution for 2 hours at room temperature, the sections were incubated ON with primary antibody solution at 4°C. Sections were then washed in PBS/0.1% Triton X-100 and incubated with the secondary antibody (1:500) and 1 mg ml^−1^ DAPI (Sigma, D9564) for 2 hours at room temperature. Details on the blocking solutions, primary and secondary antibodies used are described in the **supplementary tables T1** and **T2**. After incubation with the secondary antibodies, the sections were washed in PBS and mounted in Mowiol mounting medium.

### Imaging

The colorimetric images of SA-β-gal-stained sections were acquired using a NanoZoomer-SQ digital slide scanner (Hamamatsu) or a Leica DM2500 brightfield microscope with HC PL FLUOTAR 20x / 0.5 NA Dry objective. Immunostained sections were imaged using a motorized Zeiss Axio Observer widefield fluorescence microscope equipped with an Axiocam 506 mono CCD camera or a Zeiss Cell Observer SD confocal microscope equipped with an Evolve 512 EMCCD camera (Plan-Apochromat 20x / 0.80 NA Dry objectives). Each image is a maximum intensity projection of a z-stack acquired from the 10/12 μm cryosection. F4/80-and GAP43-stained immunosections were imaged using a Zeiss Axio Scan.Z1. The processing of acquired images was performed using Zeiss ZEN 2 (blue edition) and the image analysis software Fiji. Adobe Illustrator was used for assembly of figures.

### Quantification of SA-β-gal+ cells

To characterize the senescence profile in both models, SA-β-gal^+^ cells were manually quantified (using a Cell Counter plugin in Fiji) and averaged across 4 (zebrafish) or 8 (mouse) longitudinal sections imaged at the lesion periphery (from 0.5 to 2.5 mm laterally to the lesion) at 3, 7, 15, 30 and 60 dpi. SA-β-gal^+^ cells were quantified in the gray matter but not in the white matter and normalized to the total area covered (cells/mm^2^).

### White matter sparing

One set of sections spaced 0.1 mm apart and spanning the entire block was stained with FluoroMyelin™ Green (ThermoFisher Scientific, F34651) for 1 hour. The percentage of cross-sectional area (% CSA) with spared myelin was calculated by manually measuring the area of stained myelin in Fiji and normalizing it to the total cross-sectional area in each section (every 0.1 mm) along 2 mm rostrally and caudally from the lesion epicenter, which was identified as the section with the smallest % CSA.

### Axonal preservation

GAP43^+^ axons were counted based on previously described methods (Hata *et al.*, 2006; Almutiri *et al.*, 2018). Quantifications of GAP43^+^ fibers were performed only in the white matter in 3 longitudinal spinal sections (per biological sample) of the dorsal horn region using a custom-made macro in Fiji that, after manually establishing a threshold value and defining the lesion epicenter, determined the number of positive fibers every 1 mm from 4 mm rostral to 4 mm caudal to the lesion site and normalized it to the tissue length covered in each measurement. Axon number was calculated as a percentage towards the ratio (fibers/mm) obtained 4 mm above (rostral) the lesion, where the spinal tracts were intact.

### Fibrotic scar area

A distinct set of sections was stained with anti-PDGFRβ and anti-GFAP in order to identify the fibrotic scar area and border. The percentage of fibrotic scar area at lesion epicenter was calculated by manually outlining the PDGFRβ^+^ area and normalizing it to the total cross-sectional area. Measurements were performed using Fiji tools.

### Inflammation

Spinal sections 0.1 mm apart extending from 1 mm rostral to 1 mm caudal to the lesion epicenter were stained for the pan macrophage marker F4/80. Due to the fact that macrophages are difficult to be individually distinguished within the central lesion core, the measurements of F4/80^+^ cells were expressed as a percentage of the total cross-sectional area. The area of F4/80^+^ staining was measured using a custom-made macro in Fiji that, after manually setting a threshold value, calculated the F4/80^+^ area and normalized it to the total cross-sectional area.

### Data analysis

GraphPad Prism 7 was used for data visualization and SigmaPlot 14 for statistical analysis. The senescence profile after SCI was analyzed using a one-way ANOVA followed by a Bonferroni’s post hoc test (zebrafish) or a non-parametric Kruskal-Wallis one-way ANOVA test (mouse). BMS and Bladder Score data were analyzed using a two-way repeated-measures ANOVA, followed by a Bonferroni’s post hoc test. HL, ITP, white matter sparing, axonal preservation, fibrotic area and inflammation data were analyzed using a normal two-way ANOVA, followed by a Bonferroni’s post hoc test. All data were expressed as mean ± SEM, with statistical significance determined at *p*-values<0.05.

## ACKNOWLEDGEMENTS

We thank the support given by the Fish and Rodent Facilities, the Histology and Comparative Pathology Laboratory and the Bioimaging Unit of the Instituto de Medicina Molecular João Lobo Antunes. Ana Ribeiro and Carmen de Sena Tomás for critical reading of the manuscript.

## AUTHORS CONTRIBUTIONS

D.P.C. performed study conception and design, performed all experimental procedures in zebrafish, co-performed all experimental procedures in mice (surgeries, injuries and functional tests), performed SA-β-gal stainings and immunofluorescence, acquired and analysed data and co-wrote the manuscript. I.M. performed study design, co-performed all experimental procedures in mice (surgeries, injuries and functional tests), analysed data and co-wrote the manuscript. A.M.C. processed and cryosectioned mouse samples. A.F. processed and cryosectioned zebrafish samples. D.N.S. aided surgical procedures and perfusions. T.P. developed tools for imaging data quantifications and gave intellectual input in imaging data analysis. A.F.D. performed immunofluorescence. D.C. aided surgical procedures. A.J. performed study conception and design, supervised the project, interpreted the data. L.S. performed study conception and design, supervised the study, interpreted the data and co-wrote the manuscript.

## SUPPLEMENTARY INFORMATION

Supplementary Methods

Fig. S1. The number of SA-β-gal+ cells in Sham-injured spinal cords remains unchanged over the experimental 60 day-period.

Fig. S2. ABT-263 successfully eliminates senescent cells in the mouse spinal cord.

Fig. S3. Elimination of senescent cells with D+Q promotes motor and sensory recovery following a spinal cord injury in mice.

Fig. S4. Depletion of senescent cells leads to a reduced fibrotic scar extension following a spinal cord injury in mice.

Fig. S5. Injury biomechanics for the different experimental cohorts of C57BL/6J mice.

Table S1. List of primary antibodies.

Table S2. List of secondary antibodies.

