## Supplementary information for "Depletion of senescent cells improves functional recovery after spinal cord injury"

##### Quantification of senescence-associated β-galactosidase (SA-β-gal)$^+$ cells in Sham-injured animals

SA-β-gal$^+$ cells were manually quantified (using a Cell Counter plugin in Fiji) and averaged across 4 (zebrafish) or 8 (mouse) longitudinal sections imaged laterally to the injury segment at 3, 7, 15, 30 and 60 days post-Sham injury. SA-β-gal$^+$ cells were quantified in the gray matter but not in the white matter and normalized to the total area covered (cells/mm$^2$).

##### Quantification of SA-β-gal$^+$ cells after ABT-263 administration

The ABT-263 senolytic effect after spinal cord injury (SCI) was evaluated by manually quantifying (using a Cell Counter plugin in Fiji) and averaging the number of SA-β-gal$^+$ cells in 10 transversal sections at the lesion periphery (from 0.5 to 2.5 mm rostrally or caudally to the lesion) at 15, 30 and 60 days post-injury (dpi). SA-β-gal$^+$ cells were quantified in the gray matter but not in the white matter and normalized to the total area covered (cells/mm$^2$). Two distinct quantifications were performed: one in the total sectional grey matter and other only in the ventral horn.

##### Dasatinib + Quercetin (D+Q) administration protocol

Dasatinib (Sigma, SML2589, 5 mg/kg/day) + Quercitin (Sigma, 1592409, 50 mg/kg/day) or vehicle (10% PEG400, Sigma, 81172) were administered by oral gavage, as described previously (Zhu *et al.*, 2015), for 10 consecutive days starting at 5 dpi until 14 dpi. After spinal cord injuries, mice were randomly assigned to each group for each endpoint group, 15 dpi (n = 7) and

30 dpi (n = 15). Within the same cage animals received different treatments to exclude specific environmental cage input.

**Fibrotic scar length**

Using the set of sections stained for PDGFRβ and GFAP, the rostral and caudal extents of PDGFRβ$^+$ fibrosis were determined for each lesion, and total lesion length was calculated by multiplying the number of sections containing fibrotic tissue by the distance between each section (0.1 mm).

### Fig. S1

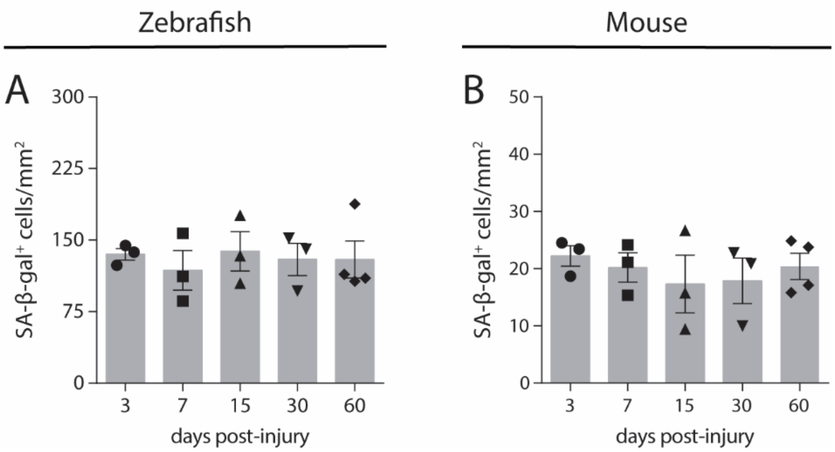

Zebrafish
Mouse
A
B
SA-β-gal⁺ cells/mm²
300
225
150
75
0
50
40
30
20
10
3
7
15
30
60
days post-injury

**Fig. S1. The number of SA-β-gal$^+$ cells in Sham-injured spinal cords remains unchanged over the experimental 60 day-period.**

SA-β-gal$^+$ cells were quantified laterally to the injury segment at 3, 7, 15, 30 and 60 days post-Sham injury. SA-β-gal$^+$ cells were quantified in the gray matter but not in the white matter and normalized to the total area covered (cells/mm$^2$). No differences were observed between the experimental end-points, meaning that, during this time window, the number of SA-β-gal$^+$ cells in the spinal cord is not affected by animal age. $n$ = 3-4. Data are presented as mean ± SEM.

#### Fig. S2

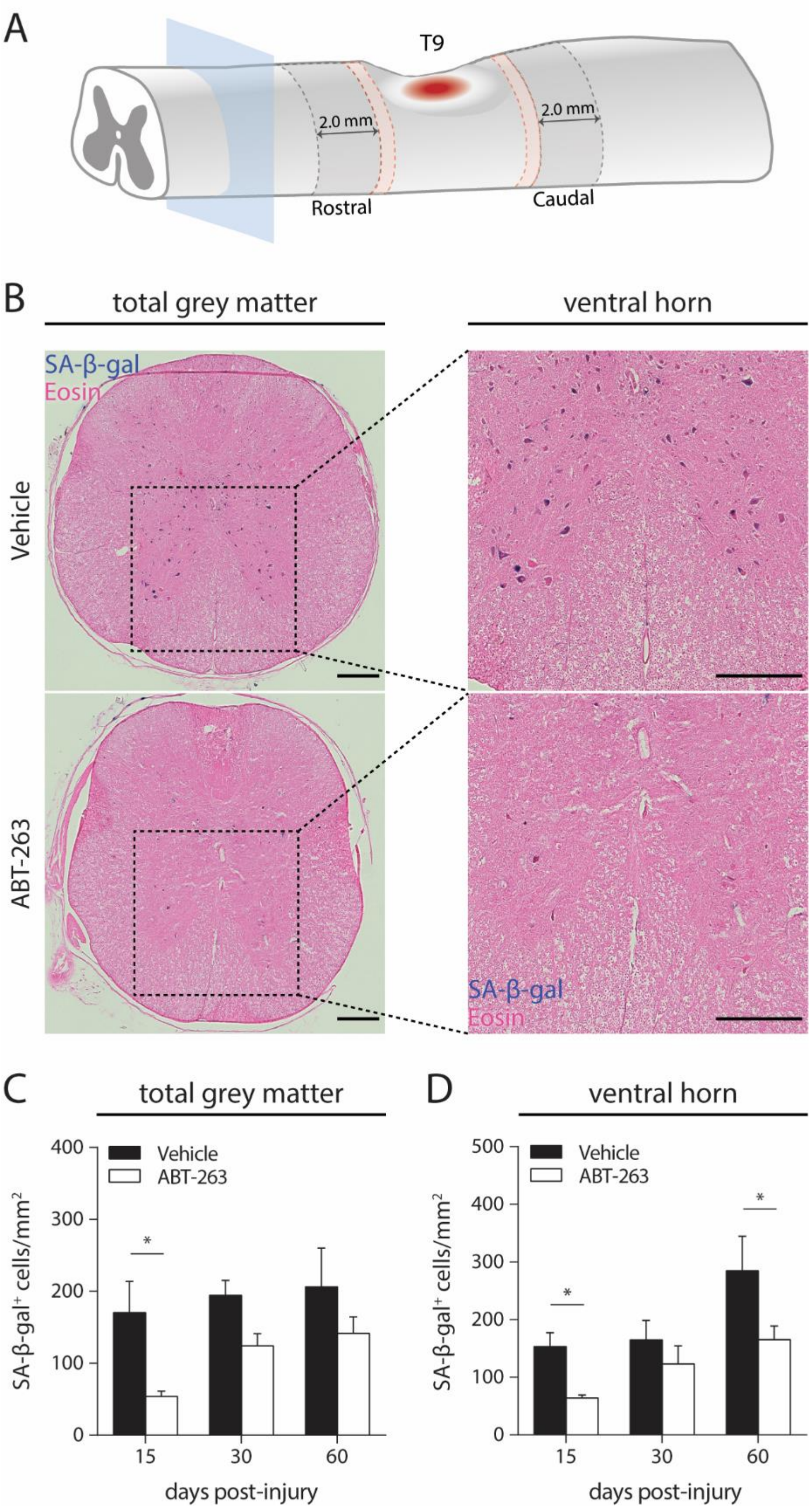

A
T9
2.0 mm
2.0 mm
Rostral
Caudal
B
total grey matter
ventral horn
SA-β-gal
Eosin
Vehicle
ABT-263
SA-β-gal
Eosin
C
total grey matter
D
ventral horn
Vehicle
ABT-263
SA-β-gal⁺ cells/mm²
days post-injury
15
30
60
0
100
200
300
400
500
*

**Fig. S2. ABT-263 successfully eliminates senescent cells in the mouse spinal cord.**

(**A**) SA-β-gal$^+$ cells were quantified in a total of 10 different transversal sections (5 rostral and 5 caudal) along 2.0 mm at the lesion periphery. A 0.5 mm interval (red dashed zone) was established between the lesion and the beginning of the quantification region. (**B**) An eosin counterstaining was performed after cryosectioning. SA-β-gal$^+$ cells (blue) were quantified in the total sectional gray matter and only at the ventral horn. Scale: 200 µm. (**C**-**D**) Quantifications were performed at all experimental endpoints (15, 30 and 60 days post-injury). At 15 dpi, ABT-263 treatment significantly decreased the number of SA-β-gal$^+$ cells/mm$^2$ in the total grey matter and in the ventral horn by 68.4% and 58.0%, respectively. At 60 dpi, a significant reduction (41.9%) of SA-β-gal$^+$ cells/mm$^2$ in ABT-263-treated animals was still observed in the ventral horn. *n* (15 dpi) = 3-4; *n* (30 dpi) = 3-4; *n* (60 dpi) = 2-3. Data are presented as mean ± SEM. **p*<0.05, ABT-263 versus Vehicle.

Fig. S3

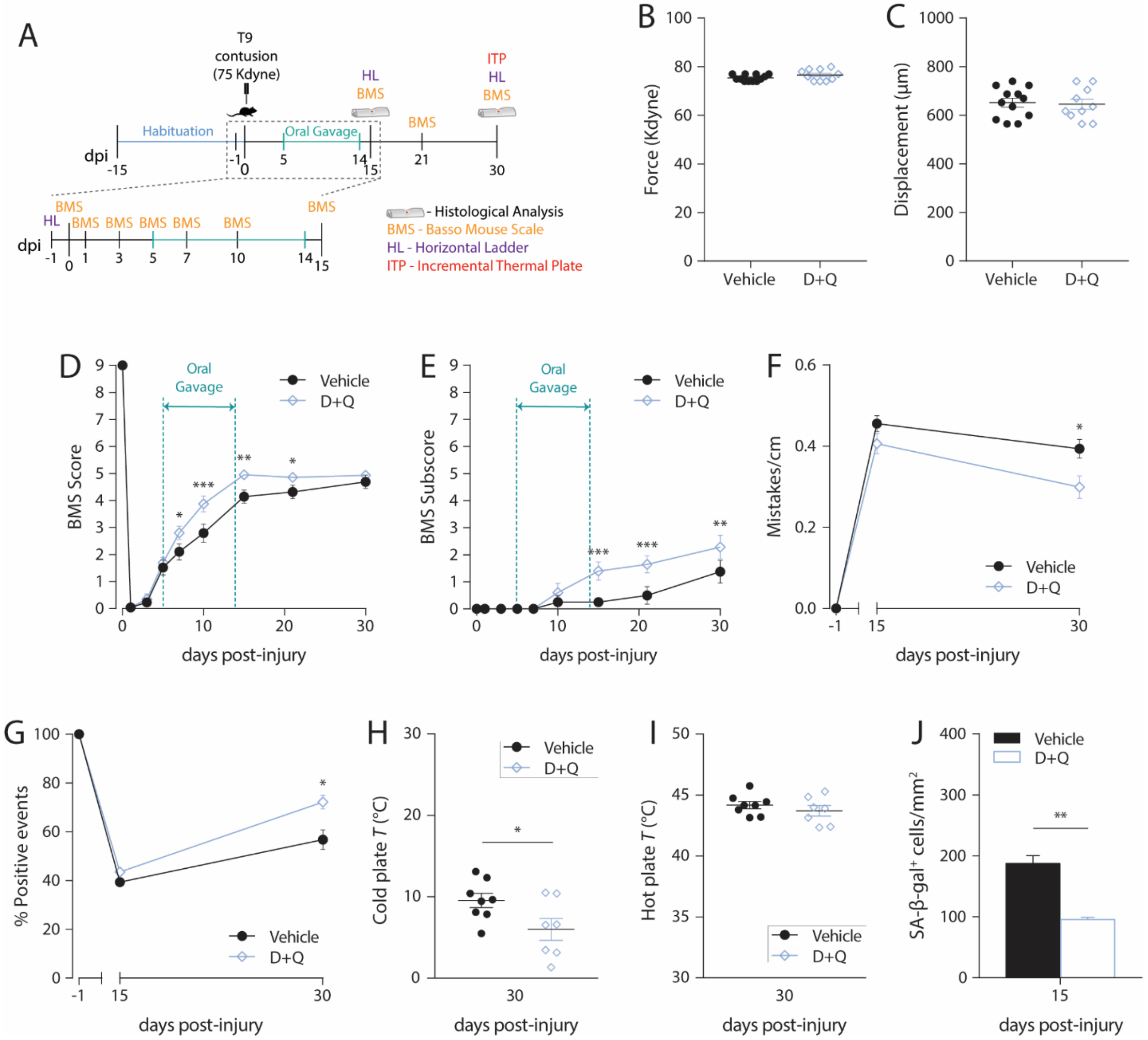
A
T9
contusion
(75 Kdyne)
HL
BMS
ITP
HL
BMS
Habituation
Oral Gavage
BMS
dpi
-15
-1
0
5
14
15
21
30
BMS
HL
BMS BMS BMS BMS BMS
BMS
dpi
-1
0
1
3
5
7
10
14
15
- Histological Analysis
BMS - Basso Mouse Scale
HL - Horizontal Ladder
ITP - Incremental Thermal Plate
B
Force (Kdyne)
Vehicle
D+Q
C
Displacement (µm)
Vehicle
D+Q
D
BMS Score
Oral
Gavage
Vehicle
D+Q
days post-injury
E
BMS Subscore
Oral
Gavage
Vehicle
D+Q
days post-injury
F
Mistakes/cm
Vehicle
D+Q
days post-injury
G
% Positive events
Vehicle
D+Q
days post-injury
H
Cold plate T (°C)
Vehicle
D+Q
days post-injury
I
Hot plate T (°C)
Vehicle
D+Q
days post-injury
J
SA-β-gal+ cells/mm2
Vehicle
D+Q
days post-injury

**Fig. S3. Elimination of senescent cells with D+Q promotes motor and sensory recovery following a spinal cord injury in mice.**

(**A**) Schematic of the experimental setup. Animals were habituated to the different behavioral setups for a 15-day period, before being submitted to a moderate-to-severe (force: 75 Kdyne; displacement: 550-750 µm) T9 contusion injury. Injured animals received daily vehicle or D+Q via oral gavage, from 5 to 14 days-post-injury (dpi). (**B**) Spinal cord impact force (in kilodynes, Kdyne) and (**C**) tissue displacement (in micrometers, µm) at the time of the contusion injury for the experimental groups. No differences were observed in force (Vehicle: 75.4 ± 0.4 Kdyne; D+Q: 76.6 ± 0.7 Kdyne) or displacement (Vehicle: 652.2 ± 18.3 µm; D+Q: 645.1 ± 20.9 µm). $n$ = 10-12. Data are presented as mean ± SEM. (**D** and **E**) Basso Mouse Locomotor Scale (BMS) score and subscore were evaluated in an open field at different time-points (0, 1, 3, 5, 7, 10, 15, 21 and 30 dpi). $n$ = 10-12. (**F** and **G**) The locomotor performance in the Horizontal Ladder (HL) was assessed at -1 (control), 15 and 30 dpi by quantifying the total number of mistakes per centimeter and the percentage of singular positive events (plantar step, toe step and skip) measured and averaged across three successful trials. $n$ = 7-10. (**H** and **I**) Thermal allodynia was tested at 30 dpi by determining the temperature at which injured mice reacted to a cold or hot stimulus. $n$ = 7-8. (**J**) At 15 dpi, SA-β-gal$^+$ cells were quantified in the total sectional gray matter in a total of 10 different transversal sections (5 rostral and 5 caudal) along 2.0 mm at the lesion periphery. D+Q treatment significantly decreased the number of SA-β-gal$^+$ cells/mm$^2$ in the total grey matter by 49.3%. $n$ = 3-4. Data are presented as mean ± SEM. $*p<0.05$, $**p<0.01$, $***p<0.001$, D+Q versus Vehicle.

#### Fig. S4

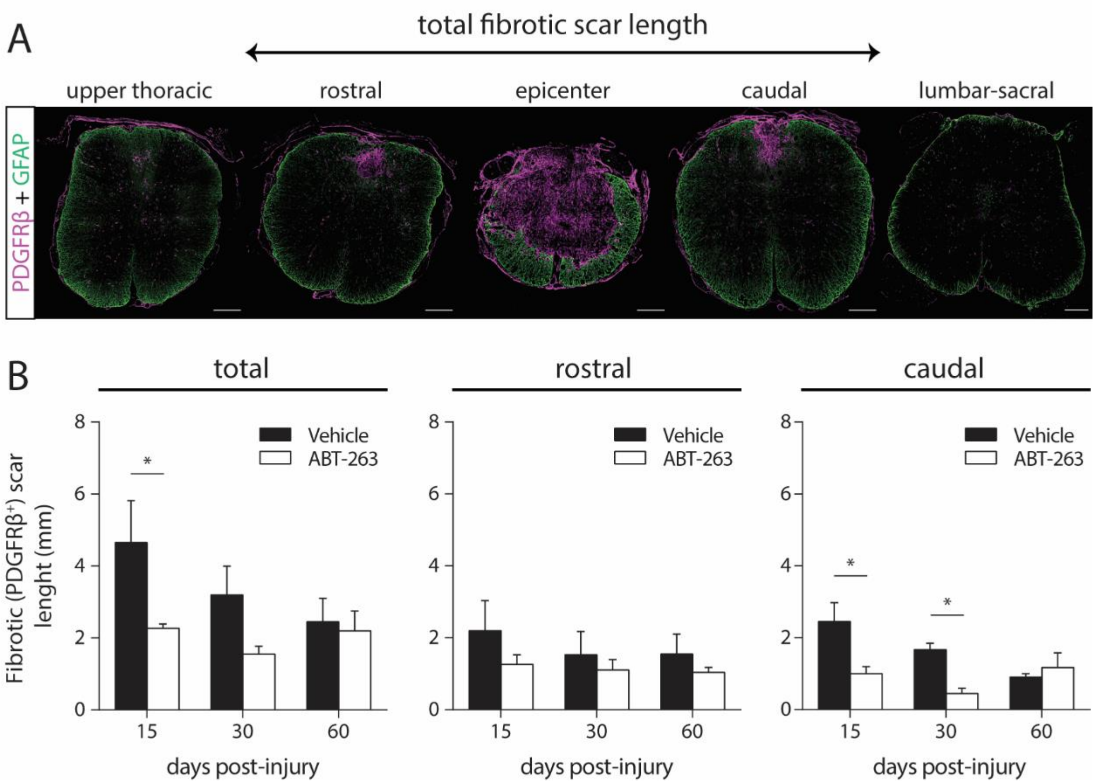
A
total fibrotic scar length
upper thoracic
rostral
epicenter
caudal
lumbar-sacral
PDGFRβ + GFAP
B
total
rostral
caudal
Vehicle
ABT-263
Fibrotic (PDGFRβ+) scar lenght (mm)
0
2
4
6
8
15
30
60
*
days post-injury

**Fig. S4. Depletion of senescent cells leads to a reduced fibrotic scar extension following a spinal cord injury in mice.**

(**A**) The total lenght of the fibrotic scar was measured by calculating the distance, rostral and caudal from the lesion epicenter, until which scarring PDGFRβ$^+$ (magenta) tissue was visualized, i.e the rostral-caudal distance spanned by sections containing fibrotic tissue. GFAP$^+$ tissue (green) surrounds the fibrotic tissue. Scale: 200 µm. (**B**) Total, rostral and caudal fibrotic extensions were calculated at 15, 30 and 60 days post-injury. *n* (15 dpi) = 3-4; *n* (30 dpi) = 3-4; *n* (60 dpi) = 2-3. Data are presented as mean ± SEM. *$p<0.05$, ABT-263 versus Vehicle.

### Fig. S5

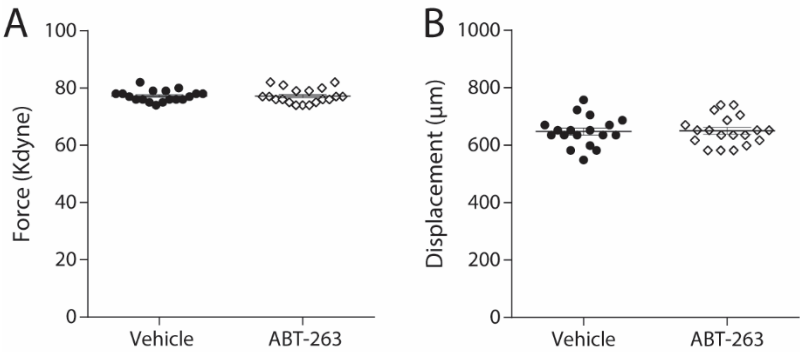
A
100
80
60
40
20
0
Force (Kdyne)
Vehicle
ABT-263
B
1000
800
600
400
200
0
Displacement (μm)
Vehicle
ABT-263

**Fig. S5. Injury biomechanics for the different experimental cohorts of C57BL/6J mice**.

(**A**) Spinal cord tissue impact force (in kilodynes, Kdyne) and (**B**) displacement (in micrometers, µm) at the time of the contusion injury for the experimental groups. No differences were observed in force (Vehicle: 77.2 ± 2.0 Kdyne; ABT-263: 77.3 ± 2.6 Kdyne) or displacement (Vehicle: 647.6 ± 12.2 µm; ABT-263: 650.4 ± 11.6 µm). *n* = 18-19. Data are presented as mean ± SEM.

**Table T1**

| Antigen | Host | Dilution | Retrieval | Blocking Solution | Source/Reference |
|---|---|---|---|---|---|
| p21 | Rabbit | 1:100 | - | 1% bovine albumin serum/1% DMSO/0.3% Tx in PBS | Santa Cruz/sc-397; RRID: AB_632126 |
| γ-H2AX | Rabbit | 1:500 | - | 1% bovine albumin serum/1% DMSO/0.3% Tx in PBS | Novus Biologicals/NB100-384; RRID: AB_10002815 |
| p16 | Rabbit | 1:50 | - | 1% bovine albumin serum/1% DMSO/0.3% Tx in PBS | ProteIntech/10883-1-AP; RRID: AB_2078303 |
| HuC/D | Mouse | 1:500 | - | 1% bovine albumin serum/0.1% Tx in PBS | Life Technologies/A21271; RRID: AB_221448 |
| NeuN | Rabbit | 1:100 | - | 5% bovine albumin serum/0.3% Tx in PBS | ProteIntech/26975-1-AP |
| GFAP | Rat | 1:400 | - | 5% goat serum/0.5% Tx in PBS | ThermoFisher Scientific/13-0300; RRID: AB_2532994 |
| PDGFRβ | Rabbit | 1:200 | - | 5% goat serum/0.5% Tx in PBS | Abcam/ab32570; RRID: AB_777165 |
| F4/80 | Rat | 1:100 | - | 5% bovine albumin serum/0.3% Tx in PBS | Abcam/ab6640; RRID: AB_1140040 |
| GAP43 | Rabbit | 1:500 | Sodium Citrate Buffer, pH 6.0 | 5% goat serum/0.3% Tx in PBS | Novus Biologicals/NB300-143; RRID: AB_10001196 |

**Table T1.** List of primary antibodies.

**Table T2**

| **Specificity** | **Host** | **Fluorophore** | **Source/Reference** |
|---|---|---|---|
| Rabbit | Goat | Alexa Fluor 488 | ThermoFisher Scientific/A11008; RRID: AB_143165 |
| Rabbit | Goat | Alexa Fluor 568 | ThermoFisher Scientific/A11011; RRID: AB_143157 |
| Rat | Goat | Alexa Fluor 488 | ThermoFisher Scientific/A11006; RRID: AB_141373 |
| Mouse | Goat | Alexa Fluor 594 | ThermoFisher Scientific/A11020; RRID: AB_141974 |

**Table T2.** List of secondary antibodies.
